# Repeated FRAP of the actin-binding protein CapG in the cell nucleus - a functional assay for EGF signaling in the single live breast cancer cell

**DOI:** 10.1101/2023.09.29.560144

**Authors:** MK Fernandez, M Sinha, M Renz

**Affiliations:** Stanford University School of Medicine

## Abstract

Compartmentalization and differential distribution of proteins within a cell maintain cellular function and viability. CapG is the only gelsolin-related actin-binding protein that distributes in steady state diffusively throughout cytoplasm and cell nucleus. To detect changes in CapG’s nuclear shuttling in response to external stimuli on the single cell level, we established repeated FRAP experiments of one and the same breast cancer cell. With this experimental set up, we found that ATP-depletion reversibly decreased CapG’s shuttling into the cell nucleus. The addition of epidermal growth factor (EGF) increased CapG’s nuclear shuttling within minutes. Serum-starvation doubled the number of breast cancer cells from 40% to 80% displaying increased CapG shuttling in response to EGF. Testing five different potential CapG phosphorylation sites, we found that serine 70 mediates the increase in CapG’s nuclear shuttling triggered by EGF. Thus, repeated FRAP of CapG in the cell nucleus can be used as functional readout of signaling cascades in the same single live breast cancer cell.

## Introduction

Compartmentalization and differential transport of solutes across membranes are critical for the living cell and determine cellular and sub-cellular functions. While the steady-state distribution of proteins can be assessed by immunohistochemistry of fixed cells, the analysis of the dynamic distribution of proteins between cellular compartments requires the use of quantitative live-cell fluorescence microscopy such as fluorescence recovery after photobleaching (FRAP)^1,2^. FRAP was established in the late 1970s to help analyze the ensemble dynamics of molecules in the plasma membrane. Photobleaching of a defined region in a cell creates spatially separated bleached and still fluorescent molecules, thus perturbs the equilibrium of fluorescent molecules and permits the subsequent monitoring of the recovery of the fluorescence equilibrium. FRAP thereby reveals the mobility and net transport of fluorescent molecules.

The gelsolin-related actin-binding protein CapG is the only member of its family that distributes in steady state diffusively and evenly throughout the entire cell, i.e., cell nucleus and cytoplasm^3^. CapG lacks the nuclear export sequence other proteins of its family comprise^4^ and shows no classic nuclear localization signal. A presumed nuclear localization sequence is of uncertain relevance^5^. It has been reported that phosphorylated CapG preferentially localizes to the cell nucleus^6^. Nuclear shuttling of CapG was shown *in vitro* to require energy and the transport protein importin β^7^. As a capping protein, CapG reversibly blocks the rapidly growing barbed ends of actin filaments^8^. The experimental overexpression or knockout of CapG has been shown to increase and decrease, respectively, the migratory potential of various cell types^9^. CapG was found to be overexpressed in cancer including breast and ovarian cancer^10^.

While the function of CapG in the cell nucleus remains unknown, it has been hypothesized that the nuclear CapG fraction is relevant for cell migration and cancer cell invasiveness^7,11^. We previously showed a difference in CapG’s nuclear shuttling in cancer and normal cells as exemplified in the breast cancer cell line MDA-MB-231 and the nearly normal breast epithelial cells MCF-12A^12^. Here, we set out to further characterize the determinants of CapG’s nucleocytoplasmic compartmentalization in the breast cancer cells MDA-MB-231 and resolve changes in CapG shuttling on the single cell level by using fluorescence recovery after photobleaching (FRAP) repeatedly in the same live cell.

## Results

Several aspects of the intracellular mobility and compartmentalization of CapG in the living cancer cell which we had characterized previously^12,13^ made us consider using repeated FRAP of the nuclear fraction of CapG as a functional single cell readout for intracellular signaling. (i) CapG displays a large mobile fraction within the cell nucleus of breast cancer cells which results in the homogeneous bleaching of the entire cell nucleus compartment even if a bleaching area is applied that is smaller than the cell nucleus. (ii) The nuclear envelope forms a significant diffusion barrier with CapG nuclear transport taking minutes, i.e., orders of magnitude longer than the bleaching time. (iii) CapG is expressed in fairly high amounts in the breast cancer cell so that repeated bleaching of the cell nucleus compartment and thereby a significant number of CapG-GFP molecules does not pose signal-to-noise problems. And (iv), the MDA-MB-231 breast cancer cells used here proved themselves resilient enough to tolerate repeated photobleaching without obvious signs of phototoxicity over many hours. Therefore, we set out to employ repeated FRAP experiments of CapG-GFP in the cell nucleus of the same live breast cancer cell as shown in **Figure 1A**. The main goal was to analyze changes in CapG-GFP’s nuclear shuttling over time in response to external stimuli on the single cell level. After a FRAP experiment was completed, we added an external stimulus, and repeated the FRAP experiment of CapG-GFP in the cell nucleus of the same cell.

**Figure 1:**
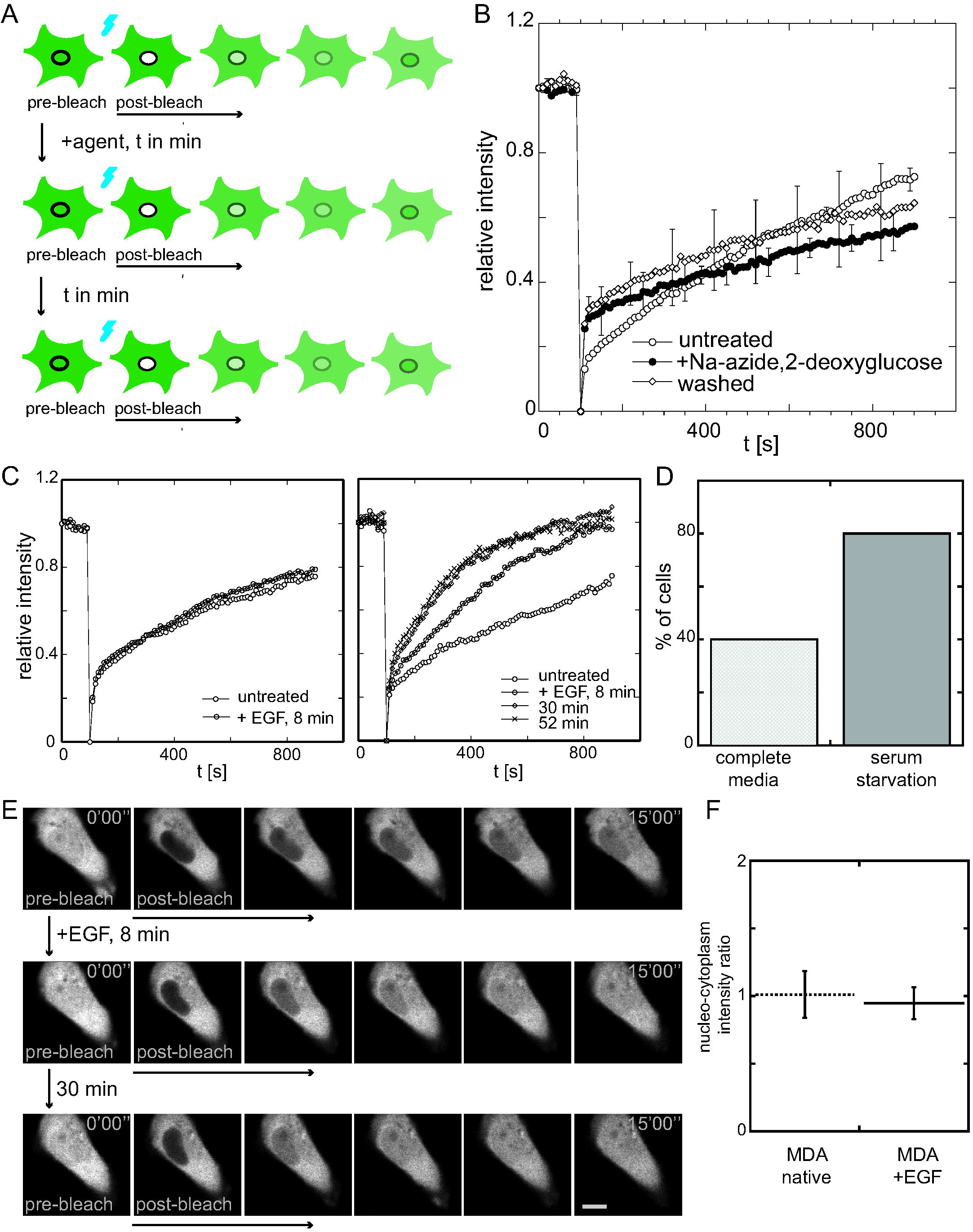
Establishing repeated FRAP of CapG in the cell nucleus of the same live breast cancer as a functional readout for intracellular signaling. **Figure 1A:** The schematic illustrates the experimental set-up of repeated FRAP of CapG in the same cell to compare nuclear shuttling before and after the addition of a stimulating or inhibiting agent. **Figure 1B:** Fluorescence recovery of CapG-GFP in MDA-MB-231 before, 15 min, and 2 h after ATP-depletion using sodium azide and 2-deoxyglucose. 3 cells were measured. Error bars indicate standard deviations. **Figure 1C:** Effect of the addition of EGF on the CapG-GFP nuclear transport in MDA-MB-231 cells. Example of a single cell recovery curve that did not show an increase in nuclear CapG-GFP shuttling, and another single cell recovery curve that did show an increase at 8 and 30 min but leveled off at 50 min. **Figure 1D:** In complete media, 4/10 (40%) MDA-MB-231 cells demonstrated increased CapG-GFP nuclear shuttling, while 8/10 (80%) MDA-MB-231 serum-starved cells displayed increased import kinetics after the addition of EGF. **Figure 1E:** MDA-MB-231 cell displaying increased nuclear CapG-GFP import after EGF addition as revealed by the repeated FRAP assay employed here. FRAP experiments performed before EGF addition, as well as 8 min and 30 min after EGF addition. Scale bar 10 um. **Figure 1F:** Steady-state distribution of CapG-GFP in single MDA-MB-231 cells before and 30 min after EGF addition. The nucleocytoplasmic fluorescence intensity ratio did not change indicating no CapG-GFP accumulation in the cell nucleus after EGF addition significant enough to be detected by confocal imaging of bulk CapG-GFP.

To first assess if our repeat FRAP assay could resolve differences in the CapG nuclear transport in the same live cancer cell, we tested its energy dependence which had been reported in *in vitro* experiments^7^. Using the described repeated FRAP assay, we added 10 mM sodium azide and 6 mM 2-deoxyglucose to achieve ATP-depletion of the cell. After 15 min, the CapG-GFP import into the cell nucleus of MDA-MB-231 cells decreased indicating an ATP-dependent transport process. The decrease in nuclear CapG-GFP import, however, was not irreversible, and cells were not driven into apoptosis, but could be rescued by washing and replacing the media with complete media (**Figure 1B**). The third FRAP measurement followed 2 h later and showed recovery and increase in nuclear CapG-GFP shuttling. The viability of the cells was not compromised, neither by the temporary ATP-depletion nor the repeated photobleaching.

Next, we tested if the repeated FRAP assay could be used to analyze the effect of external stimuli on CapG’s nuclear shuttling. The epidermal growth factor (EGF) is a known tumor promoting factor and has been used as chemoattractant in migration assays. EGF receptors are expressed in MDA-MB-231 and known to increase their migratory potential^14^. Since the nuclear fraction of CapG has been hypothesized to be critical for cancer cell invasion^7^, we set out to analyze the effect of EGF on the CapG distribution in the live cell. We hypothesized that EGF may increase nuclear CapG shuttling. After measuring the baseline CapG-GFP transport kinetics in a cell, we added EGF to an end concentration of 200 ng/mL. Because we anticipated a fast non-genomic effect, we repeated the FRAP measurements in the same cell 8 min, 30 min, and 50 min after the addition of EGF. Some MDA-MB-231 cells did not display any change in fluorescence recovery kinetics, other cells however did (**Figure 1C**). For the cells that responded to EGF, the steepness of the relative fluorescence intensity curve increased after 8 min and further after 30 min but began to level off at 50 min as shown in **Figure 1C**. Fitting a single exponential function to the recovery curves shown in **Figure 1C** revealed a characteristic recovery time of 731.7 ± 60.9 s before the addition of EGF, 547.8 ± 21.3 s 8 min after EGF, 283.5 ± 5.7 s at 30 min and 226.6 ± 5.8 s at 50 min after EGF addition. Thus, in this representative cell the nuclear transport of CapG-GFP increased 3.2-fold over 50 min (**Figure 1E, Supplemental Movie 1**). Analyzing 10 MDA-MB-231 cells, 4/10 (40%) cells showed an increase in CapG-GFP nuclear transport 30 min after the EGF addition, while six cells did not. The differing responses of the cancer cells may reflect distinct cell cycle states or indicate that the EGF signaling cascade is not expressed or not active in every cell. To assess a possible cell-cycle dependent EGF response, we serum-starved cells for 20 h and thereby arrested them in the G0 phase. Under these conditions, the addition of EGF triggered in 8/10 (80%) of the analyzed cells an increase in CapG-GFP nuclear shuttling (**Figure 1D**). The addition of EGF did not shift the steady-state distribution of CapG-GFP significantly enough to be detected by repeat confocal imaging, neither in MDA-MB-231 cells grown in complete media nor in the serum-starved cells. Confocal images were taken to measure the nucleocytoplasmic fluorescence intensity ratio in steady state before and 30 min after the addition of EGF (**Figure 1F**). No accumulation of CapG-GFP in the cell nucleus was noted in steady state bulk imaging despite the increased nuclear transport as measured by FRAP.

We then tested which phosphorylation sites in the CapG amino acid sequence are critical for the increase in EGF stimulated CapG nuclear shuttling. To narrow down potential candidates, we used an online available phosphorylation site prediction tool (NetPhos-3.1, DTU Health Tech). In the literature, serines S10 and S337 as well as threonine T212 have been reported to be potential phosphorylation sites^15^. Furthermore, serine S337 has been hypothesized to be located within a protein kinase C site recognition motif^16^. We replaced the serines S10 and S337 and the threonine T212 with alanines. To increase the chance of detecting accelerated CapG-GFP nuclear import after the addition of EGF, all of the following repeated FRAP experiments were performed with serum-starved MDA-MB-231 cells. In cells expressing the triple CapG-GFP mutant *S10A, *T212A, and *S337A, an increase in CapG-GFP nuclear import was noted after the addition of EGF (**Figure 2B**). We also tested the amino acid point mutation *S337A in isolation, which again demonstrated stimulation of the CapG-GFP nuclear import by EGF (**Figure 2C**). The CapG mutant *S70A and *S200A, however, did not show any increase in CapG-GFP nuclear transport in the 10 cells analyzed (**Figure 2D**). Cells expressing the CapG-GFP mutant with the *S200A mutation alone demonstrated an increase in CapG-GFP nuclear shuttling (**Figure 2E**), while cells expressing the CapG-GFP mutant with the isolated point mutation *S70A did not (**Figure 2F**). Based on the presented repeated FRAP experiments, serine S70 is the critical CapG phosphorylation site whose phosphorylation in response to EGF signaling results in an increase in nuclear transport of CapG (**Figure 3**).

**Figure 2:**
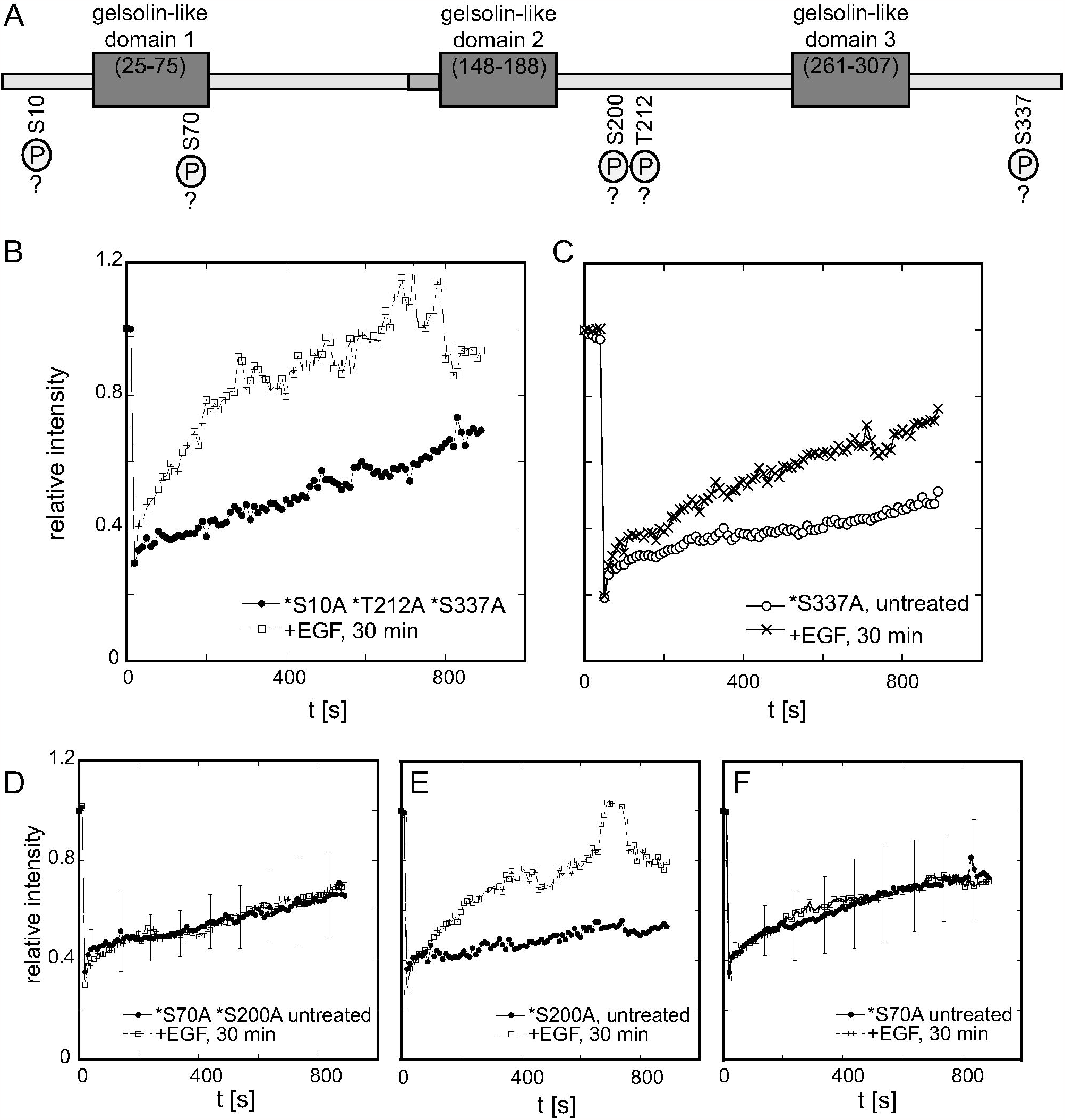
Screening for posttranslational modifications after EGF stimulation using repeated FRAP of CapG in the cell nucleus of the same live breast cancer reveals critical CapG phosphorylation site. **Figure 2A:** Schematic of the CapG protein sequence with its 3 gelsolin-like repeats. Potential phosphorylation sites that were analyzed in this study are outlined: S10, S70, S200, T212, and S337. **Figure 2B:** Serum-starved MDA-MB-231 cell expressing the CapG-GFP mutant *S10A, *T212A, *S337A displayed a steeper fluorescence recovery curve 30 min after EGF addition. **Figure 2C:** Serum-starved MDA-MB-231 cell expressing the CapG-GFP mutant *S337A displayed a steeper fluorescence recovery curve 30 min after EGF addition. **Figure 2D:** 0/10 (0%) serum-starved MDA-MB-231 cells expressing the CapG-GFP mutant *S70A, *S200A displayed increased fluorescence recovery kinetics 30 min after EGF addition. Error bars indicate standard deviations. **Figure 2E:** Serum-starved MDA-MB-231 cell expressing the CapG-GFP mutant *S200A displayed a steeper fluorescence recovery curve 30 min after EGF addition. **Figure 2F:** 0/10 (0%) serum-starved MDA-MB-231 cells expressing the CapG-GFP mutant *S70A displayed increased fluorescence recovery kinetics 30 min after EGF addition. Error bars indicate standard deviations.

**Figure 3:**
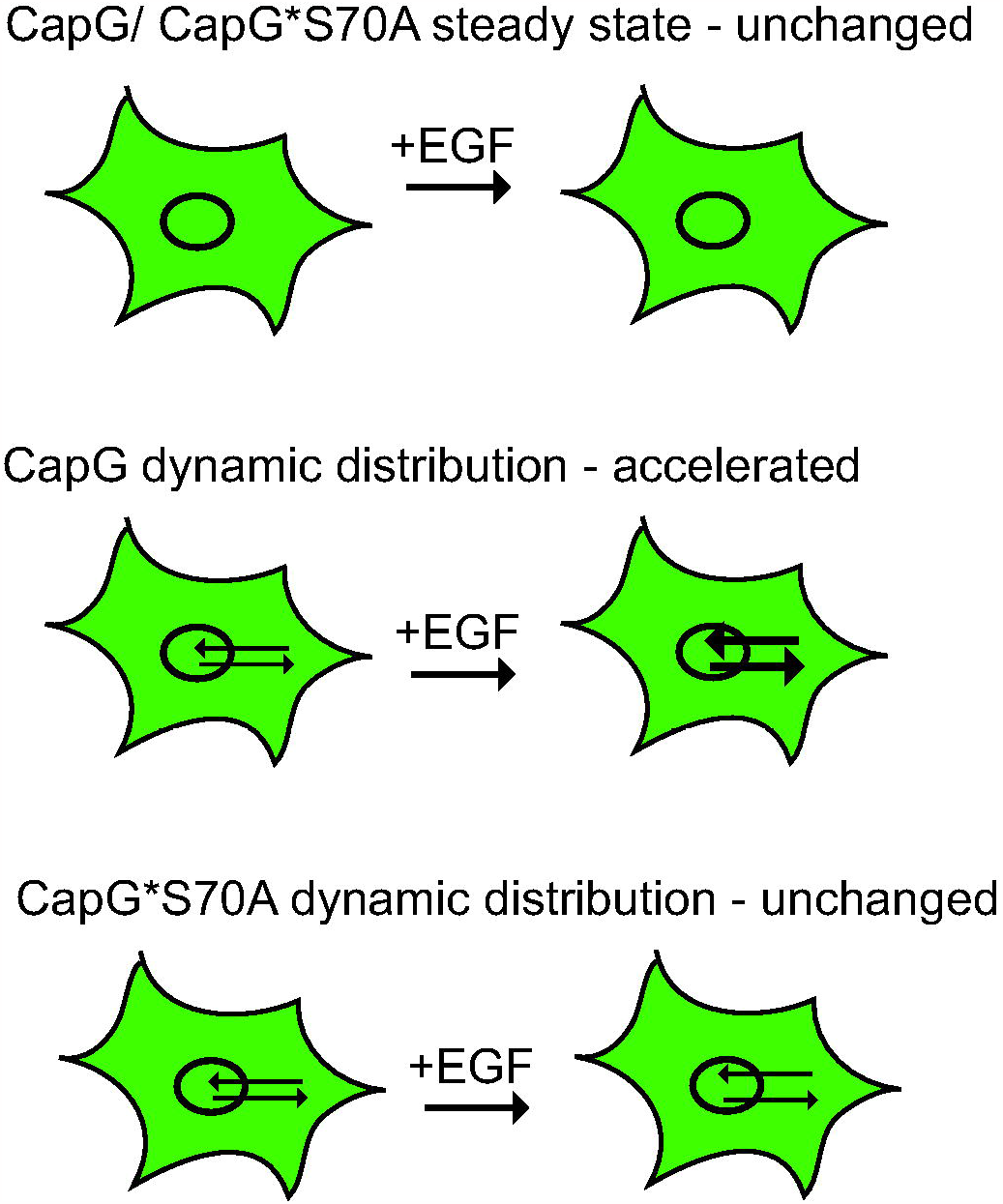
Model schematic of CapG distribution and nucleocytoplasmic shuttling in the live cell. While no changes are detectable in steady state, CapG nucleocytoplasmic shuttling is accelerated after EGF addition. Given the lack of significant CapG-GFP accumulation in the cell nucleus, this likely reflects an increase in im- and export of CapG. Serine S70 is the critical CapG phosphorylation site downstream of the EGF signaling cascade.

## Discussion

We showed that EGF increases nuclear shuttling of CapG in the breast cancer cells MDA-MB-231 within minutes. The critical phosphorylation site for this increase is the serine S70 based on our FRAP experiments. Serine S70 phosphorylation may increase nuclear CapG shuttling by different mechanisms including (i) increased binding to importin β or other shuttle proteins and (ii) inducing conformational change that decreases CapG’s hydrodynamic radius and thereby facilitates CapG transport. Repeated FRAP of CapG in the cancer cell nucleus permits the assessment of the functionality of signaling cascades on the single cell level provided those signaling cascades affect CapG shuttling. A functional single cell assay seems especially important since upstream receptors of signaling cascades may not be expressed in every cell or may not be biologically active even if expressed which has been reported for the EGF receptor in breast cancer cells^17^ and ensemble analyses may average out and thus miss single-cell behavior. In our hands, repeated FRAP experiments in the same live cell did not trigger any apparent cell toxicity and cells could be imaged within any signs of compromise over many hours.

The noted increase in CapG cell nuclear import is likely paralleled by an increased CapG export since no significant accumulation of CapG in the cell nucleus was noted on the ensemble or bulk level. Our previous FRAP analyses suggested an only small CapG fraction of 3% that is bound and immobilized in the cell nucleus of MDA-MB-231 cells^13^. The biological meaning of the noted increased nucleocytoplasmic CapG shuttling is thus far uncertain and relates to the open question of CapG’s function in the cell nucleus. Various binding partners in the cell nucleus have been suggested in the literature and include steroid receptors^5,18^, NF-κB^19^, and/or nuclear actin.

The high abundance of the freely diffusive CapG in the cell nucleus may serve as a pool of capping proteins for transient need, as it were ready on demand. Calculations of CapG in the cytoplasm showed a 5 to 10-fold abundance relative to the plus ends of actin filaments in macrophages^20^. It is also feasible that CapG co-shuttles and transports cargo into the cell nucleus and that such cargo in turn exerts the biological function in the cell nucleus. Studies addressing these questions are ongoing.

In summary, we showed that repeated FRAP of CapG in the cell nucleus is feasible and can resolve changes in nuclear shuttling which are inaccessible to repeated static imaging.

Repeated FRAP of CapG in the cell nucleus is a suitable functional assay to assess downstream effects of signaling cascades in the single living cancer cell and demonstrated that CapG’s nuclear shuttling increased within minutes after EGF stimulation and depends on the phosphorylation of serine S70.

## Material and Methods

### Plasmids and transfections

We used the plasmid CapG-eGFP as previously published^12^. For simplicity, we call the construct CapG-GFP throughout the manuscript. Site-directed mutagenesis was performed using the QuikChange Kit from Agilent according to the manufacturer’s protocol. The serines S10, S70, S200, S337 and the threonine at T212 were replaced by alanines. The following primers were used 5’-ATT CCC CAG AGT GGC GCT CCA TTC CAG GCT CAG T-3’ for *S10A, 5’-CCA GCA GTC AGC CCG GGA TGA GCA -3’ for *S70A, 5’-GCC ATC CGG GAC GCG GAG CGA CAG GGC AA-3’ for *S200A, 5’-CAG GTG GAG ATT GTC GCT GAT GGG GAG GAG CCT GCT-3’ for *T212A, 5’-TCT GCC TCA GGG CCG TGA GGC TCC CAT CTT CAA GCA ATT T-3’ for *S337A, respectively. MDA-MB-231 cells were transfected with the CapG-GFP construct and its mutants using Transfectin (Biorad, Hercules, CA) according to the manufacturer’s protocol.

### Cell culture

The MDA-MB-231 cell line, a human breast cancer cell line, was obtained from ATCC (HTB-26) and cultured in DMEM without phenol red and supplemented with 10% fetal bovine serum (FBS) and 1% glutamic acid.

### FRAP and confocal microscopy

The Zeiss laser-scanning confocal microscope LSM710 with a 25-mW Argon ion laser and a 62x 1.4 N.A. objective was used to perform the photobleaching experiments. The FRAP experiment was performed by exposing defined regions of cells to 100% laser intensity for 20 iterations. Images were taken with laser intensity set to 0.1% as indicated by the control software.

For quantitative analysis, regions were defined on the acquired images delineating the bleached nucleus, the cytoplasm and the extracellular space. By integrating over all pixels using the image processing software Fiji (NIH, Bethesda), the fluorescence intensity was determined for each time point in each region. The determined mean gray values were background subtracted, corrected for acquisition photobleaching, laser intensity fluctuations and loss of fluorescence due to the bleaching event. The data were normalized to the initial fluorescence intensity. The following equation for the normalized fluorescence was applied:

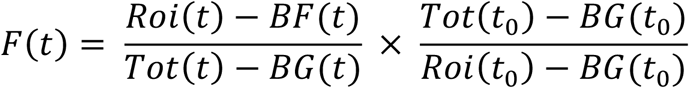

where Roi(t) denotes the fluorescence intensity at time t in the region of interest, i.e., the cell nucleus, Tot(t) denotes the fluorescence intensity in the cytoplasm at each time point and BG(t) the extracellular background fluorescence in each time point. The normalized relative fluorescence intensity curve was fit with the single exponential function *F* (*t*)= 1 − [a − *b*(1 − *e*^− *λt*^], where *a* is the fraction of fluorescence intensity initially bleached, b the fraction that recovered after the time *t* with the rate *λ*. The recovery time τ is the reciprocal of the recovery rate *λ*.

FRAP experiments were performed at 37°C in Hepes-buffered medium (pH 7.3) 20 h after transfection. For serum starvation, cells were washed with Hanks balanced salt solution, and DMEM media with 0.1% fetal calf serum and 1% Glutamine were added to the cells. Cells were incubated in this media 20 h before the experiments. Epidermal growth factor (EGF, Sigma) was reconstituted as per manufacturer’s instructions. A 100 ug/ mL stock solution was aliquoted and stored at -80°C. EGF was added for the experiments to the cell culture media in an end concentration of 200 ng/mL.

## Author contributions

MKF, MS and MR performed the experiments. MR analyzed the data and wrote the manuscript. MKF and MS edited the manuscript.

## Acknowledgements

We would like to thank the core facility Neuroscience Microscopy Service at Stanford University for the use of the Zeiss LSM 710.

## Conflict of interest statement

The authors have no conflict of interest.

## Figure legends

**Supplemental Movie:** MDA-MB-231 cell displays increased nuclear CapG-GFP import after EGF addition as revealed by the repeat FRAP assay of the same cell.

## Notes

### Competing Interest Statement

The authors have declared no competing interest.

## References

1. Axelrod D, Koppel DE, Schlessinger J, Elson E, Webb WW. Mobility measurement by analysis of fluorescence photobleaching recovery kinetics. Biophys J. 1976;16(9):1055–1069.

2. Rabut G, Ellenberg J. Photobleaching techniques to study mobility and molecular dynamics of proteins in live cells: FRAP, iFRAP, and FLIP. In: Goldman R, Spector DL, eds. Live cell imaging. A laboratory manual. Cold Spring Harbor: Cold Spring Harbor Laboratory Press; 2005:101–126.

3. Prendergast GC, Ziff EB. Mbh 1: a novel gelsolin/severin-related protein which binds actin in vitro and exhibits nuclear localization in vivo. EMBO J. 1991;10(4):757–766.

4. Van Impe K, De Corte V, Eichinger L, Bruyneel E, Mareel M, Vandekerckhove J, Gettemans J. The Nucleo-cytoplasmic actin-binding protein CapG lacks a nuclear export sequence present in structurally related proteins. J Biol Chem. 2003;278(20):17945–17952.

5. Gettemans J, Van Impe K, Delanote V, Hubert T, Vandekerckhove J, De Corte V. Nuclear actin-binding proteins as modulators of gene transcription. Traffic. 2005;6(10):847–857.

6. Onoda K, Yin HL. gCap39 is phosphorylated. Stimulation by okadaic acid and preferential association with nuclei. J Biol Chem. 1993;268(6):4106–4112.

7. De Corte V, Van Impe K, Bruyneel E, Boucherie C, Mareel M, Vandekerckhove J, Gettemans J. Increased importin-beta-dependent nuclear import of the actin modulating protein CapG promotes cell invasion. J Cell Sci. 2004;117(Pt 22):5283–5292.

8. Southwick FS, DiNubile MJ. Rabbit alveolar macrophages contain a Ca2+-sensitive, 41,000-dalton protein which reversibly blocks the “barbed” ends of actin filaments but does not sever them. J Biol Chem. 1986;261(30):14191–14195.

9. Thompson CC, Ashcroft FJ, Patel S, et al. Pancreatic cancer cells overexpress gelsolin family-capping proteins, which contribute to their cell motility. Gut. 2007;56(1):95–106.

10. Dahl E, Sadr-Nabavi A, Klopocki E, et al. Systematic identification and molecular characterization of genes differentially expressed in breast and ovarian cancer. J Pathol. 2005;205(1):21–28.

11. Pellieux C, Desgeorges A, Pigeon CH, Chambaz C, Yin H, Hayoz D, Silacci P. Cap G, a gelsolin family protein modulating protective effects of unidirectional shear stress. J Biol Chem. 2003;278(31):29136–29144.

12. Renz M, Betz B, Niederacher D, Bender HG, Langowski J. Invasive breast cancer cells exhibit increased mobility of the actin-binding protein CapG. Int J Cancer. 2008;122(7):1476–1482.

13. Renz M, Langowski J. Dynamics of the CapG actin-binding protein in the cell nucleus studied by FRAP and FCS. Chromosome Res. 2008;16(3):427–437.

14. Price JT, Tiganis T, Agarwal A, Djakiew D, Thompson EW. Epidermal growth factor promotes MDA-MB-231 breast cancer cell migration through a phosphatidylinositol 3’-kinase and phospholipase C-dependent mechanism. Cancer Res. 1999;59(21):5475–5478.

15. Papala A, Sylvester M, Dyballa-Rukes N, Metzger S, D’Haese J. Isolation and characterization of human CapG expressed and post-translationally modified in Pichia pastoris. Protein Expr Purif. 2017;134:25–37.

16. Glaser J, Neumann MH, Mei Q, et al. Macrophage capping protein CapG is a putative oncogene involved in migration and invasiveness in ovarian carcinoma. Biomed Res Int. 2014;2014:379847.

17. Ali R, Wendt MK. The paradoxical functions of EGFR during breast cancer progression. Signal Transduct Target Ther. 2017;2:16042–.

18. Lee YH, Campbell HD, Stallcup MR. Developmentally essential protein flightless I is a nuclear receptor coactivator with actin binding activity. Mol Cell Biol. 2004;24(5):2103–2117.

19. Ma Q, Zhao M, Long B, Li H. Super-enhancer-associated gene CAPG promotes AML progression. Commun Biol. 2023;6(1):622.

20. Young CL, Feierstein A, Southwick FS. Calcium regulation of actin filament capping and monomer binding by macrophage capping protein. J Biol Chem. 1994;269(19):13997–14002.

